# Bayesian surprise tracks the strength of perceptual insight

**DOI:** 10.64898/2026.02.26.708200

**Authors:** Johannah Völler, Juan Linde-Domingo, Carlos González-García

## Abstract

Suddenly finding the solution to a problem after a period of impasse often comes with a feeling of insight. This subjective experience is proposed to arise as a consequence of prediction errors. Accordingly, previous studies have revealed that more incorrect initial predictions result in more intense insights. Crucially however, prominent models of Bayesian inference suggest levels of computationally-defined surprise are not a simple feature of distance between predictions and inputs, but also their precision or certainty. Yet, how these two factors interact to give rise to insight experiences remains unknown. In this pre-registered study, participants were exposed to ambiguous images while they tried to guess the correct label of the image (to derive prediction accuracy) and rated their confidence in that label (for prediction uncertainty). We then measured the intensity of their insight when a solution was given. As predicted, we found that the intensity of insight was a result of both the prediction accuracy and the uncertainty awarded to it. More specifically, when initial predictions were far from the true label, those made with lower confidence induced weaker insights, while the opposite pattern was observed when predictions were closer to the reality. Trial-by-trial estimations of prediction errors from participants’ responses closely mirrored insight ratings. Finally, we analysed data from two additional independent datasets with different modalities and setups and replicated the interaction between prediction accuracy and uncertainty on the intensity of insight. Altogether, these findings suggest that insight experiences are read out from prediction errors and highlight the key role of uncertainty in characterising this relationship.

## Introduction

To make sense of an inherently noisy world, the brain iteratively generates and tests predictions about the causes underlying the sensory input (Clark, 2013; Friston, 2005). Incorporating prior knowledge following such a recurrent architecture helps reduce ambiguity efficiently in most scenarios, i.e., the mismatch (prediction error) between the brain’s initial guess and the incoming evidence can be resolved rapidly (Lee & Mumford, 2003). In some cases, however, prior knowledge fails or even impedes to resolve prediction errors, leading to a stage of impasse. For instance, in the famous nine-dot-problem, in which people have to connect nine dots arranged in a square using four straight lines, prior knowledge constrains solvers to assume the lines must stay within the square’s boundaries (Öllinger et al., 2014). Successful problem solving involves relaxing those constraints and realising that trajectories must extend beyond the boundaries.

When a solution is suddenly found after such an impasse via gaining new or restructuring preexisting knowledge, an insight can arise (Becker et al., 2025; Wiley & Danek, 2024). A frequent marker of insights is the “Aha!” experience which accompanies the solution with feelings of happiness, surprise, and confidence (Becker & Cabeza, 2025; Danek et al., 2014a; Danek & Wiley, 2017). Notably, this subjective experience is associated with more correct responses (Danek et al., 2014b; Salvi et al., 2016) and better memory performance (Becker et al., 2025; Danek et al., 2013; Kizilirmak et al., 2016), demonstrating its adaptive function in guiding learning and memory. Current theories have conceptualized this feeling of insight as a result of the sudden, larger than expected reduction in prediction error once a solution is found (Becker & Cabeza, 2025; Friston et al., 2017; Laukkonen et al., 2023). Consistent with this explanation, studies have repeatedly reported a positive link between the intensity of insight experiences and the magnitude of the prediction error (Dubey et al., 2023; Savinova & Korovkin, 2022; Van de Cruys et al., 2021). These studies have mostly inferred prediction errors from the accuracy of the initial prediction, e.g., higher Aha! for more incorrect predictions regarding the solvability of the problem (Becker et al., 2024).

However, prediction errors are not only determined by accuracy but are also strongly influenced by the (un)certainty of the initial prediction. In current models of Bayesian inference, this uncertainty, or precision, plays a key role in weighting predictions and sensory evidence based on their respective noise, giving more weight to reliable sources (Fleming, 2024a; Hohwy, 2012; Yon & Frith, 2021). Specifically, if a prediction is made with high uncertainty (low precision), deviations from it are not as surprising. Conversely, if a prediction is more certain (precise), small deviations can signal important changes (Press et al., 2020; Yon & Frith, 2021). We do not know however how these computations determine subjective insight. Consider again the nine-dot problem. If people had a strong expectation that the line has to stay inside the square’s boundaries, they might be more surprised once the solution is discovered compared to solvers who did not have such strong expectations. This uncertainty-weighting is captured within mathematical notations of Bayesian inference, where predictions and incoming evidence are two probability distributions over a hypothesis space with a mean tendency and a given variance (Ma et al., 2023). The prediction error (or Bayesian surprise) can be operationalised with the *Kullback-Leibler Divergence* (Kullback, 1959), calculating the non-overlap between the belief about the world before (prior) and after (posterior) the incoming evidence is incorporated (Friston, 2010; Press et al., 2020). More specifically, the lesser their overlap, resulting from the distance in mean tendency between the distributions and their respective variances, the higher the surprise (Itti & Baldi, 2009). Consequently, if the prediction is highly accurate, a high (low) precision will lead to very little (strong) surprise, while a high (low) precision would lead to stronger (lesser) surprise if the prediction was inaccurate (Fig. 1c). Despite the importance of considering both the prediction accuracy and its precision to describe prediction errors (Hohwy, 2012), these two characteristics have not been studied together in explaining insight experiences.

**Figure 1.**
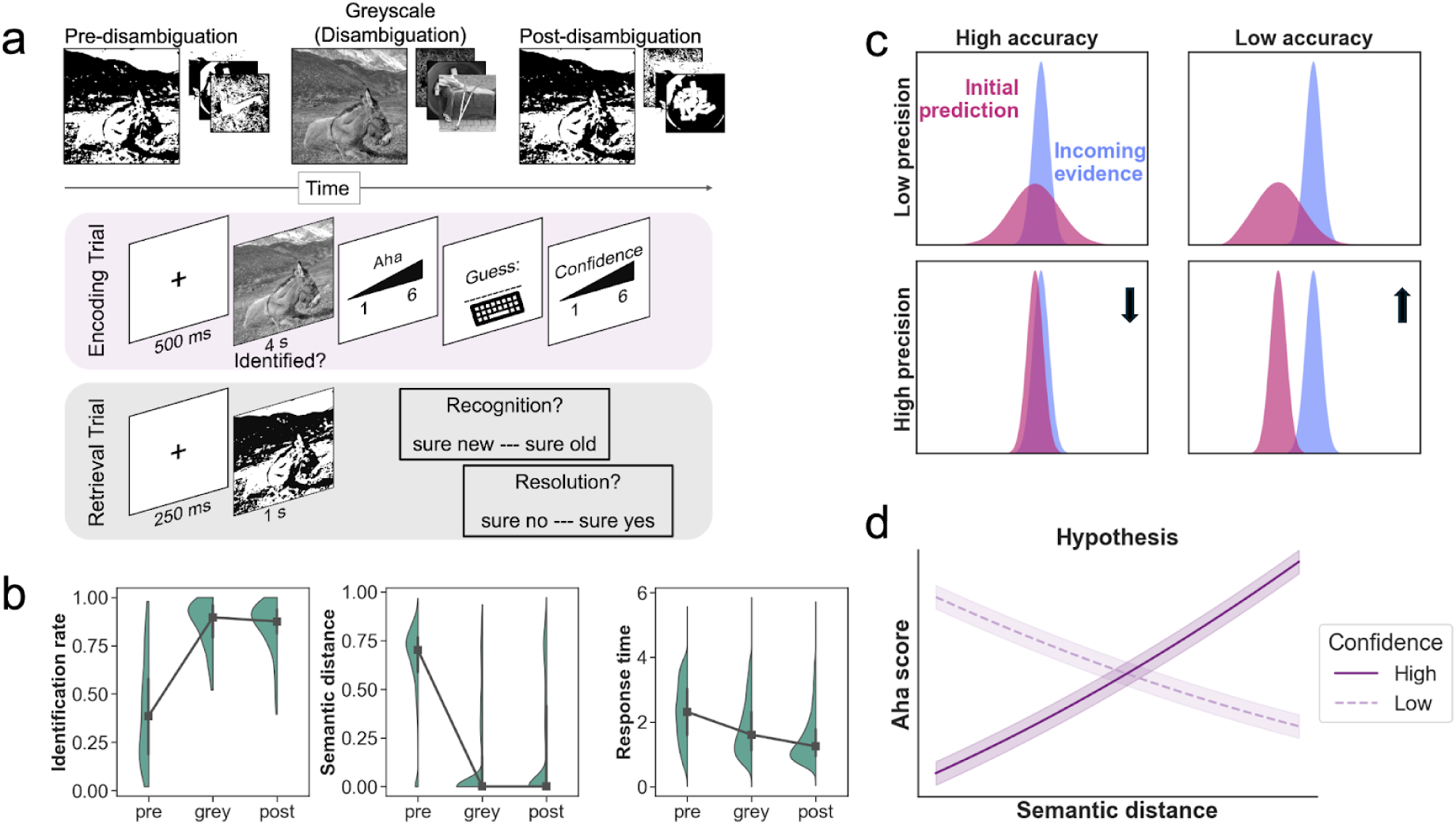
Experimental procedure, the effect of viewing stage, and hypothesis. **a**. A total of 122 participants viewed 48 different Mooney images in three stages: pre-disambiguation, clear greyscale image (disambiguation), and post-disambiguation. Images were presented in an interleaved design, such that up to 10 trials could appear in between the Mooney and its corresponding greyscale image. For every image, participants reported subjective identification via pressing the spacebar while the image was on the screen, the intensity of their Aha! (if subjectively identified), provided a typed label (guess) of the main object in the image, and rated their confidence in the label. In a surprise immediate retrieval test, participants rated their recognition memory strength and whether they subjectively resolved (identified) the Mooney image on 6-point likert scales for half of the encoded images and the same amount of new Mooney images. Mooney and clear images in the figure are extracted from THINGS-Mooney (Linde-Domingo et al., 2024), which is based on THINGSplus (Stoinski et al., 2023), and are copyright-free. **b**. Being exposed to the clear greyscale image facilitated the identification of the Mooney images at post-disambiguation, subjectively (left) and also objectively as measured in semantic distance (middle). Response times (s) decreased linearly over the different viewing stages (right). Black squares represent medians, black lines denote the interquartile range, and green shaded areas indicate the data distribution. **c**. According to Bayesian inference, prediction errors are characterized not only by the accuracy of the prediction but also by their precision (inverse variance). For a given incoming evidence and a highly accurate prediction (left column), the prediction error is attenuated if the prediction is very precise, i.e., the initial prediction and the incoming evidence largely overlap. If the prediction is not accurate (right column), the resulting prediction error will be stronger for very precise predictions in comparison to imprecise predictions, as predicted by the lesser overlap between the two distributions (bottom right). Probability density functions (y-axis) are plotted over a created linespace (x-axis; −10, 15, 1000). **d**. Our hypothesis regarding the intensity of insight follows this same intuition: Aha! scores will be higher for highly accurate but imprecise guesses (left-hand side), and for inaccurate but precise guesses (right-hand side).

In this preregistered study (see Methods), we formally test the idea that the intensity of insight experiences is not exclusively explained by the accuracy of initial predictions, but also by the uncertainty awarded to them, following Bayesian inference accounts (Fig. 1d). More specifically, we expected low confidence (i.e., low precision) to result in stronger insight for lower semantic distance (higher prediction accuracy), while high confidence should result in stronger insight for higher semantic distance (lower prediction accuracy). Participants viewed ambiguous two-tone images, so called Mooney images (Mooney, 1957; Fig. 1a), for which they tried to guess the correct label and reported their confidence in that guess. Thus, guesses were defined by their distance to the correct label (prediction accuracy) and by the confidence awarded to them (prediction precision). Subsequently, participants reported the intensity of their insight once the clear version of the Mooney image (Fig. 1a) was presented, which suddenly makes the images easily identifiable (Ishikawa & Mogi, 2011; Suzuki et al., 2018).

In line with our hypothesis, we observed that initial predictions with higher semantic distance from the true label resulted in stronger insight upon disambiguation. Crucially, when initial predictions were far from the correct label, higher confidence increased the intensity of insight, while the effect was reversed for fairly correct initial guesses. We replicated this interaction in two additional independent datasets (N = 30 and N = 107) with different data collection modalities (online vs. in-lab) and conditions (0% vs. 25% vs. 50% catch trials; see below). Estimated prediction errors from semantic distance and confidence ratings using Kullback-Leibler Divergence paralleled this interaction effect, which we found to be weaker in lower precision environments. Lastly, in a surprise memory test for half of the encoded ambiguous images, we replicated previous findings linking stronger insight to improved subsequent memory (Becker et al., 2025; Danek et al., 2013; Kizilirmak et al., 2016). Altogether, these results reveal a strong parallelism between prediction error computation and insight experiences and confirm the crucial role of uncertainty in determining their intensity.

## Results

### Perceptual insight increases task performance and memory

Mooney images were presented in three viewing stages: pre-disambiguation, greyscale image (clear image to induce disambiguation), and post-disambiguation (Fig. 1a; see Methods for a more detailed description). For every image at every stage, 122 participants reported subjective identification, the intensity of their insight (rating the Aha! for subjectively identified images), and provided a typed label of the Mooney image’s main object. They were then asked to rate their confidence in the provided label. Prediction accuracy was operationalised continuously as the distance in semantic space between the participant-provided label and the target label at pre-disambiguation (see Methods). Thus, when the provided label (e.g., car) was semantically distant from the target (e.g., donkey), prediction accuracy was low. Prediction precision was approximated through participants’ confidence ratings on the provided verbal labels at pre-disambiguation, with low scores indicating more uncertain guesses as a result of imprecise available evidence (Fleming, 2024b; Geurts et al., 2022).

In line with previous research (Flounders et al., 2019; González-García et al., 2018; Linde-Domingo et al., 2025), being exposed to the clear greyscale image drastically improved performance from pre- to post-disambiguation in several behavioral indices (Fig. 1b). Repeated-measure ANOVAs on averaged data per subject and condition revealed a main effect of viewing stage on subjective identification (F_(2,242)_ = 401.36, p < 0.001, ηp^2^ = 0.77), where identification rate was significantly lower during pre-disambiguation (M = 0.41, SD = 0.49) compared to greyscale (M = 0.86, SD = 0.35; t_(121)_ = 20.21, p < 0.001, ηp^2^ = 0.56) and post-disambiguation (M = 0.85, SD = 0.36; t_(121)_ = 21.82, p < 0.001, ηp^2^ = 0.54). Similarly, semantic distance, i.e., the cosine dissimilarity between the embeddings of the provided label and the true one, was significantly higher in pre-disambiguation trials (M = 0.63, SD = 0.22) compared to greyscale (M = 0.12, SD = 0.23; t_(121)_ = −76.14, p < 0.001, ηp^2^ = 0.92) and post-disambiguation (M = 0.19, SD = 0.3; t_(121)_ = −58.23, p < 0.001, ηp^2^ = 0.87), reflected in a main effect of viewing stage (F_(2,242)_ = 3738.47, p < 0.001, ηp^2^ = 0.97). Specifically, while only 8.11% (SD = 27%) of pre-disambiguation labels were correct (i.e., semantic distance, including synonyms, was zero), naming accuracy increased to 65.93% (SD = 47%) at post-disambiguation. Lastly, response times decreased linearly over viewing stages (F_(2,242)_ = 408.02, p < 0.001, ηp^2^ = 0.77; grey-pre: t_(121)_ = −16.71, p < 0.001, ηp^2^ = 0.31; grey-post: t_(121)_ = 13.84, p < 0.001, ηp^2^ = 0.14; post-pre: t_(121)_ = −24.74, p < 0.001, ηp^2^ = 0.56), from pre-(M = 2.30, SD = 0.91) to greyscale (M = 1.79, SD = 0.84) to post-disambiguation (M = 1.46, SD = 0.72).

Next, before testing the effect of participants’ predictions on the intensity of their insight experiences, we aimed to confirm that the Aha! scores captured in our task reproduced previously reported patterns. To do so, linear mixed models were fitted to the recognition memory (sure new - sure old) and subjective resolution scores (sure no - sure yes) of Mooney images obtained in a surprise immediate retrieval test. Memory was predicted from Aha! ratings during greyscale viewing. Models included correctly disambiguated trials only to isolate the effect of insight, while spontaneous disambiguation, i.e., correctly named pre-disambiguation trials, were excluded (Ludmer et al., 2011; Van de Cruys et al., 2021). As reported before (Becker et al., 2025; Danek et al., 2013; Kizilirmak et al., 2016), Aha! scores significantly and positively predicted both recognition memory strength (χ^2^_(3)_ = 18.37, p < 0.001, B = 0.07, 95% CI [0.01, 0.12]) and subjective resolution (χ^2^_(3)_ = 23.64, p < 0.001, B = 0.07, 95% CI [0.01, 0.13]), confirming that the Aha! ratings measured in our task are behaviourally meaningful markers of the subjective experience of insight.

### Semantic distance predicts the intensity of insight experiences

To test the effect of semantic distance (as proxy for prediction accuracy) on the intensity of insight upon disambiguation (greyscale stage), we ran linear mixed models, entering only correctly disambiguated trials and subjective identification as a covariate of no interest (see Methods). Correct disambiguation was defined as post-disambiguation semantic distance of zero (including synonyms). Mixed models revealed a main effect of semantic distance on Aha! scores (χ^2^ = 269.53, p < 0.001, likelihood ratio test), with higher semantic distance leading to stronger Aha! scores upon correct disambiguation, independent of whether the image was subjectively identified at pre-disambiguation (B = 0.07, 95% CI [0.03, 0.11]). Following recent suggestions that pop-out solutions (< 2 seconds) are not considered insight solutions (Becker et al., 2021, 2024), we reran the analysis excluding those trials. Semantic distance continued to have a main effect on Aha! (χ^2^ = 87.48, p < 0.001) in the same direction (B = 0.06, 95% CI [0.02, 0.09]). We confirmed all analysis with cumulative link mixed models to account for the ordinal nature of the data (see Supplementary Material).

### Confidence moderates the effect of semantic distance on insight

Next, we tested the role of confidence (as a proxy for prediction precision) on this relationship between semantic distance and Aha! scores. Again, we only considered correctly disambiguated trials. As predicted, mixed models revealed a significant interaction between semantic distance and confidence (χ^2^_(1)_ = 14.75, p < 0.001, B = 0.04, 95% CI [0.02, 0.06]; Fig. 2a, 2b). Model comparison showed that the interaction model provided a better fit to the data (AIC = 10718.14), outperforming the semantic distance only model (AIC = 10736.72). Over the full range of both predictors, confidence had a significant negative slope for low semantic distance, but a positive slope for high semantic distance, indicating that confidence reduced the intensity of insight when the guess was fairly correct, but increased it when the guess was far from the correct label (see dashed vertical lines, Johnson-Neyman interval [−0.03, 0.43]; FDR-corrected interval [−0.05, 0.49]; Fig. 2a). Simple slope analysis indicated that this interaction was specifically driven by a significant positive slope of semantic distance at high levels (+ 1 SD) of confidence (b = 0.52, t_(3462)_ = 3.76, p < 0.001, 95% CI [0.25, 0.79]), while the semantic distance of mean confidence (b = 0.25, t_(3462)_ = 1.77, p = 0.08, 95% CI [−0.03, 0.52]) and low confidence (−1 SD) guesses was not significantly related to Aha! scores (b = −0.02, t_(3462)_ = −0.13, p = 0.90, 95% CI [−0.36, 0.32]). Again, the interaction effect was not dependent on pop-out solutions trials (χ^2^_(1)_ = 12.82, p < 0.001, B = 0.04, 95% CI [0.02, 0.06]) or on subjective identification at pre-disambiguation.

**Figure 2.**
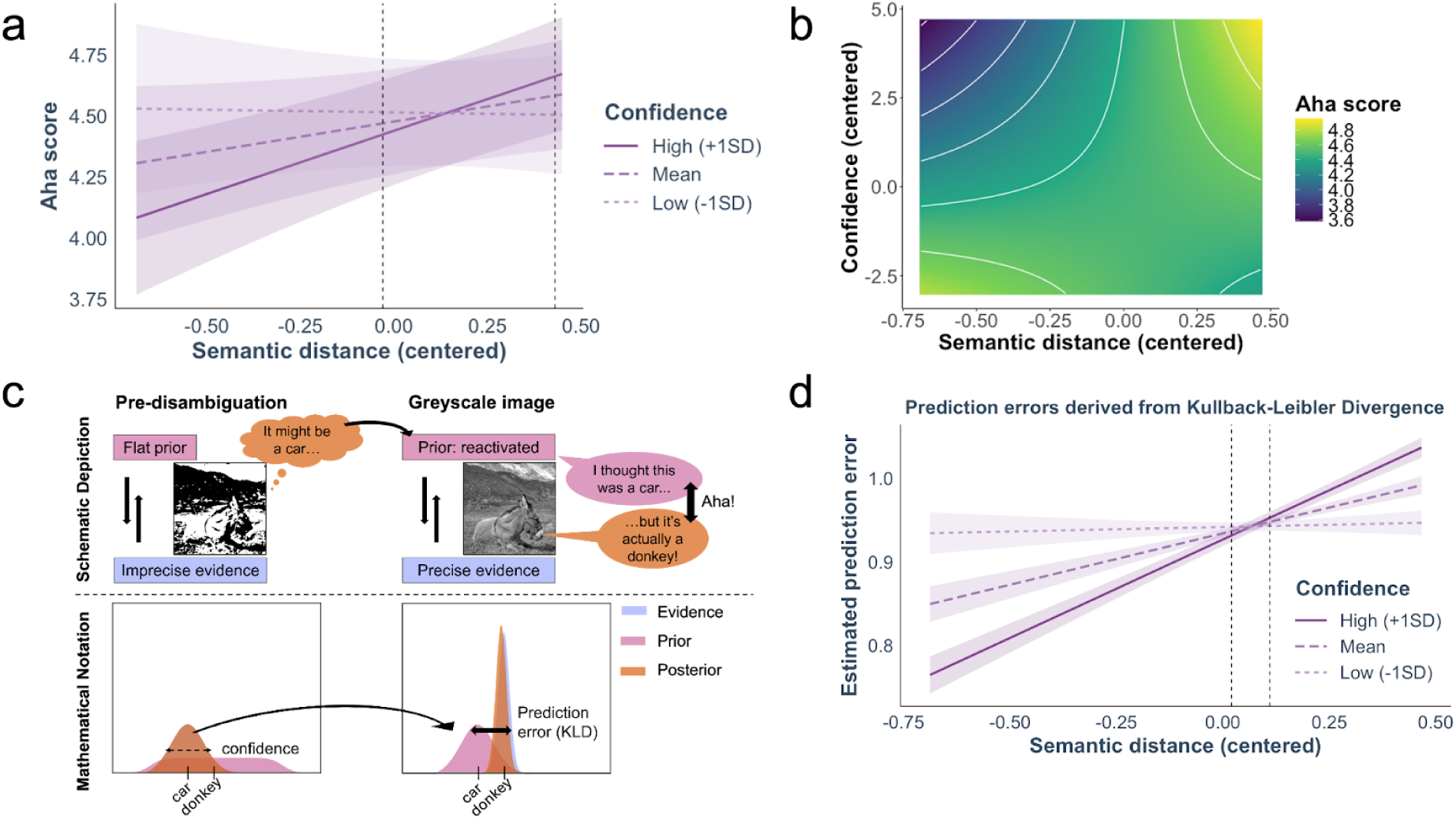
Confidence moderates the effect of semantic distance on the intensity of insight. **a**. Model-predicted Aha! values for high (+ 1SD), mean, and low (−1 SD) levels of confidence indicate that high confidence reduces Aha! ratings for low semantic distance guesses, but increases them for (very) high semantic distance guesses. Shaded areas represent 95% confidence intervals, and dashed vertical lines denote the Johnson-Neyman interval (uncorrected) outside of which confidence has a significant slope. **b**. Depicts the model-predicted Aha! values over the full range of both predictors. As can be seen, higher Aha! scores are reported for low semantic distance - low confidence and high semantic distance - high confidence guesses. White contour lines represent constant values (iso-levels) of model-predicted Aha! values, overlaid on the heatmap to highlight regions of similar predicted intensity. **c**. Estimating prediction errors using Kullback-Leibler Divergence (KLD). At pre-disambiguation, participants’ flat prior is iteratively combined with imprecise evidence (the ambiguous Mooney images), resulting in a posterior that resembles the imprecise evidence (“It might be a car…”). Mathematically, participants behavioural reports, i.e., their provided guess and the confidence rating, are read out from the posterior, such as that the posterior’s mean is defined as the semantic distance between the provided verbal label and the correct label, while the confidence rating reflects the posterior’s precision. During greyscale image viewing, the posterior from pre-disambiguation is reactivated (“I thought this was a car…”) and acts as a new prior. Iteratively combined with the highly precise evidence, the prior is updated accordingly (“…but it’s actually a donkey!”), and the ubiquitous insight is experienced. Mathematically, the KLD is calculated between the prior (posterior from pre-disambiguation) and the current posterior as an estimate for the size of the prediction error, which is then convolved with a small error term sampled from a Gaussian distribution. Please note that the evidence in both cases is displayed underneath the posterior. **d**. The KLD-derived prediction errors show the same interaction effect between semantic distance and confidence. As before, shaded areas represent 95% confidence intervals, and dashed vertical lines denote the Johnson-Neyman interval (uncorrected) outside of which confidence has a significant slope.

### Estimated trial-by-trial prediction errors behave similarly as Aha! ratings

Although our results produced the expected pattern, they are agnostic regarding the extent to which the subjective experience of insight in this task is a readout of actual prediction errors. To explore this question, we aimed at deriving prediction errors from our behavioral variables and then examined if they followed a similar interaction as the Aha! scores. An approach that has been proposed in the past to approximate prediction errors consists in calculating the Kullback-Leibler Divergence between the prior and the posterior (prior multiplied with the sensory evidence), in our case, when exposed to unambiguous, clear images (Itti & Baldi, 2009; Press et al., 2020). To do so, one needs to define the sensory evidence distribution (likelihood) and the prior. During clear image viewing, the sensory evidence represents the ground truth of the image. We thus fixed the likelihood’s mean to 0 (such as that the prior’s mean reflected the distance to a correct verbal label with a semantic distance of 0) and the variance to 0.1, to account for the fact that the sensory input at that stage (clear image) is very precise, although not of a 100% precision due to inherent noise in the sensory system (Barlow, 1956; Neri, 2010).

Next, the prior needs to be defined. Priors are often established experimentally by manipulating conditional probabilities of event-outcome associations, thereby inducing different levels of prediction errors (e.g., Ortiz-Tudela et al., 2023; Thomas et al., 2024). Here, however, the source of the prediction (and its error) is internal to the participant, making it more difficult to quantify. Crucially, in our paradigm, participants reported a typed guess and the related confidence regarding the content of the image during pre-disambiguation trials. As the posterior reflects the stabilised percept, we reasoned that participant’s responses in pre-disambiguation trials are read out from this posterior, consistent with frequent proposals of confidence ratings as a marker of the posterior’s precision (Geurts et al., 2022; Pouget et al., 2016). Thus, we treated the initial semantic distance and the confidence ratings as markers of the posterior’s mean and variance in pre-disambiguation trials, respectively. When presented with the corresponding clear version of that image, the pre-disambiguation posterior now acts as a new prior, reflecting the inherent iterative characteristics of Bayesian inference (cf. Ma et al., 2023). Following this rationale, we derived trial-by-trial prediction errors by computing the Kullback-Leibler Divergence between the prior (μ = pre-disambiguation semantic distance, σ = pre-disambiguation confidence rating) and the posterior (the prior convolved with the likelihood: μ = 0, σ = 0.1) while viewing clear, unambiguous images (Fig. 2c). Finally, to mimic inherent distortions in participant’s own readouts, prediction errors were estimated by adding a small error term sampled from a Gaussian distribution (μ = 0, σ = 0.1) to the raw Kulback-Leiber Divergence values.

We first confirmed that derived prediction errors tracked the reported intensity of insight in the task by fitting a linear mixed model, predicting Aha! scores from derived prediction errors. Indeed, prediction errors significantly predicted Aha! scores (χ^2^_(1)_ = 31.47, p < 0.001, B = 0.07, 95% CI [0.04, 0.09]), indicating that larger prediction errors led to stronger reported insight on a trial-by-trial level. The derived prediction error values were then analysed in the same way as the Aha! scores. Linear mixed models revealed a significant interaction between semantic distance and confidence on the derived prediction error values (χ^2^_(1)_ = 258.46, p < 0.001, B = 0.22, 95% CI [0.19, 0.24]; Fig. 2d). Again, confidence had a significant negative slope for low semantic distance, but a significant positive slope for high semantic distance guesses (see dashed vertical lines, Johnson-Neyman interval [0.02, 0.11]; FDR-corrected interval [0.02, 0.11]; Fig. 2d). Such reversal of the effect suggests that higher confidence resulted in weaker prediction errors for fairly correct guesses, but in stronger prediction errors for semantically distant guesses. The effect was driven by a significant positive slope of semantic distance at high (+ 1 SD; b = 0.24, t_(3833)_ = 15.53, p < 0.001, 95% CI [0.21, 0.27]) and mean levels of confidence (b = 0.12, t_(3833)_ = 9.03, p < 0.001, 95% CI [0.10, 0.15]), while low confident guesses (−1 SD) were not significantly related to prediction error values (b = 0.01, t_(3833)_ = 0.64, p = 0.52, 95% CI [−0.02, 0.04]).

### The interaction between semantic distance and confidence replicates in two independent datasets

To confirm the robustness of this finding, we analysed the relationship between semantic distance and confidence in two other datasets that were collected on two independent samples for different purposes but included the variables of interest. The “Catch-Lab” dataset was collected as part of an in-lab study with a final sample of 30 native Spanish participants. The “Catch-Online” dataset was collected online with a total of 107 English speaking participants. Both datasets included catch trials, which are trials in which a different, non-corresponding clear image was presented in-between the two Mooney image presentations to prevent disambiguation (“Catch-Lab”: 25% catch, 75% regular trials; “Catch-Online”: 50% catch, 50% regular trials), on top of “regular” trials that induce disambiguation as in the main dataset. In both datasets, confidence and Aha! was rated on visual analogue scales. Again, we only analysed regular and correctly disambiguated trials (as a control, catch trials were analysed separately).

In the “Catch-Lab” dataset, mixed models revealed a significant interaction between semantic distance and confidence (χ^2^ = 8.38, p = 0.004, B = 0.09, 95% CI [0.03, 0.16]; Fig. 3a, 3b) but no main effect of semantic distance (B = 0.08, 95% CI [−0.03, 0.20]). Over the full range of values, confidence had a significant negative slope for low semantic distance, while a significant positive slope was observed for high semantic distance (see dashed vertical line, Johnson-Neymann interval [−0.41, 0.10]; FDR-corrected interval [−0.85, 0.18]; Fig. 3a). Thus, higher confidence resulted in less intense insights for initial guesses that were close to the correct label, but this pattern reversed for semantically distant guesses. Again, the interaction effect was specifically driven by a strong positive slope of semantic distance for high confidence guesses (b = 23.93, t_(583)_ = 2.94, p = 0.01, 95% CI [7.96, 39.89]), with no significant slope for mean (b = 11.37, t_(583)_ = 1.43, p = 0.16, 95% CI [−4.18, 26.93]) and low confidence (b = −1.18, t_(583)_ = −0.12, p = 0.91, 95% CI [−20.45, 18.10]) guesses.

**Figure 3.**
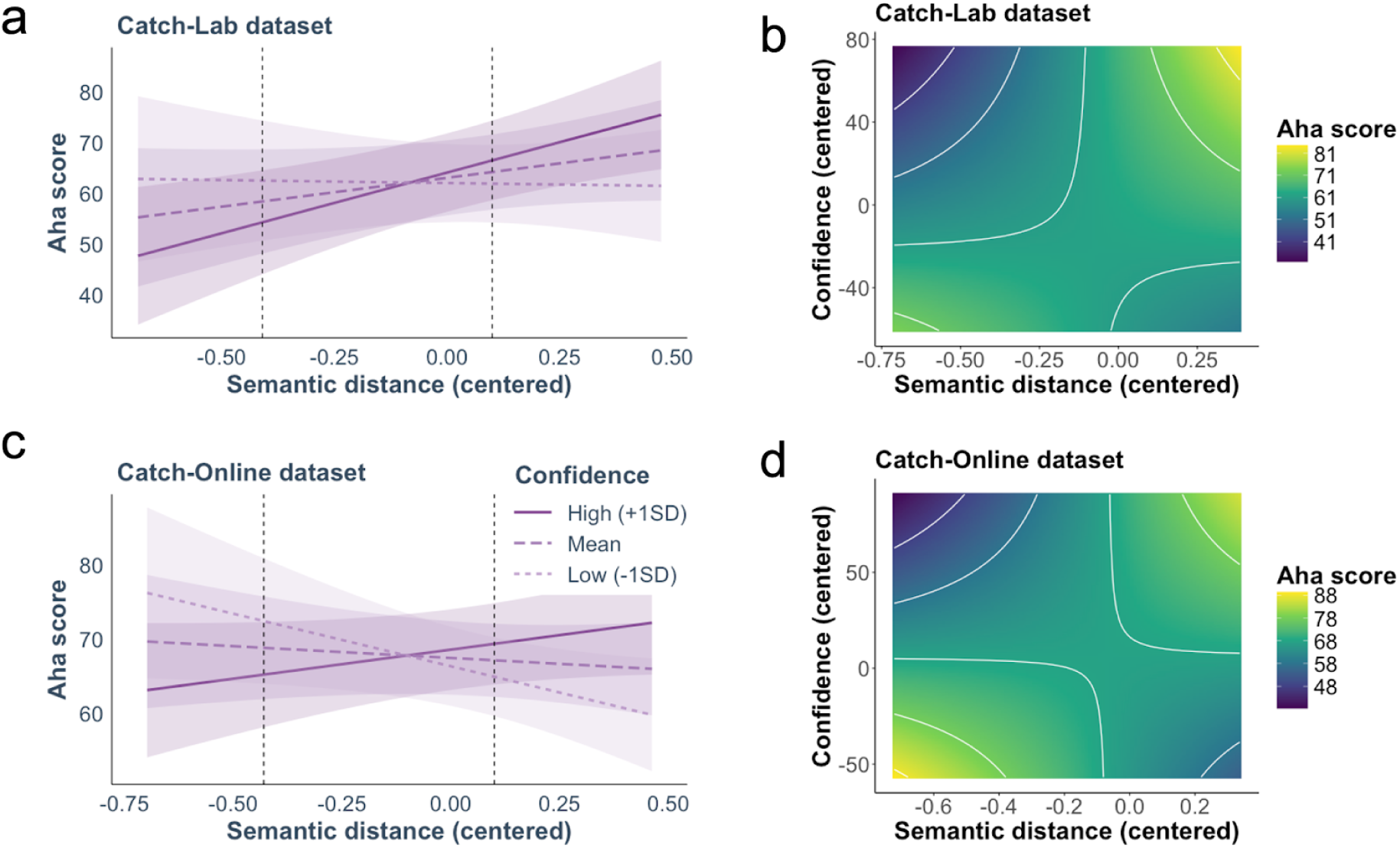
Replication of the semantic distance and confidence interaction in two independent datasets. **a**. In the “Catch-Lab” dataset, higher confidence reduced Aha! scores for low semantic distance, but increased them for high semantic distance guesses. Shaded areas represent 95% confidence intervals, and dashed vertical lines denote the Johnson-Neyman interval (uncorrected) outside of which confidence has a significant slope. **b**. Heatmap of the model-predicted values of the insight experiences in the “Catch-Lab” dataset over the full value range of both predictors. White contour lines represent iso-levels of model-predicted Aha! scores. **c**. In the “Catch-Online” dataset, higher confidence again showed a reversed effect: for low semantic distance guesses, high confidence resulted in less intense insights, but in more intense insights for higher semantic distance. Shaded areas represent 95% confidence intervals, and dashed vertical lines denote the Johnson-Neyman interval (uncorrected) outside of which confidence has a significant slope. **d**. Heatmap of the model-predicted Aha! values in the “Catch-Online” dataset over the full value range of both predictors. White contour lines represent iso-levels of model-predicted Aha! values.

Similarly, there was a significant interaction between semantic distance and confidence in the “Catch-Online” dataset (χ^2^_(1)_ = 9.84, p = 0.002, B = 0.08, 95% CI [0.03, 0.12]; Fig. 3c, 3d) but no main effect of semantic distance (B = −0.02, 95% CI [−0.09, 0.05]). Considering the full range of both regressors, higher confidence significantly reduced Aha! ratings for low semantic distance guesses, and increased Aha! ratings for high semantic distance guesses (see dashed vertical lines, Johnson-Neyman interval [−0.43, 0.10]; FDR-corrected interval [−0.72, 0.18]; Fig. 3c). This time, simple slope analysis revealed no significant slope of semantic distance on Aha! scores for high (b = 7.85, t_(734)_ = 1.53, p = 0.13, 95% CI [−2.19, 17.88]) and mean confidence guesses (b = −3.18, t_(734)_ = −0.63, p = 0.53, 95% CI [−13.02, 6.66]). Instead, the interaction was the result of a significant negative slope of semantic distance on Aha! scores for low confident guesses (b = −14.20, t_(734)_ = −2.05, p = 0.04, 95% CI [−27.76, −0.65]).

Crucially, when these analyses were performed in catch trials that control for the mere effect of image repetition, there was no significant interaction effect in either of the two datasets. Specifically, in the “Catch-Lab” dataset, the model containing the interaction between semantic distance and confidence was not significantly better than the model containing both independent variables but not their interaction (χ^2^_(1)_ = 1.85, p = 0.17, B = −0.04, 95% CI [−0.10, 0.02]). Likewise, the interaction model in the “Catch-Online” dataset was not significantly better than the baseline model (χ^2^_(1)_ = 0.41, p = 0.52, B = −0.01, 95% CI [−0.04, 0.02]). This indicates that the observed interaction is specific to resolving the ambiguity through connecting the Mooney image with its corresponding clear version, and not a mere effect of perceiving a clear image.

### Low precision environments attenuate the effect of confidence

So far, the results reveal a consistent effect of confidence about our own predictions on subsequent insight experiences. However, other sources of uncertainty, such as the sensory input, could influence this relationship. Similar to the precision of the prediction, the input precision modulates prediction errors, such as that deviations from a given prediction should be less surprising if the sensory environment is more variable or volatile (Yon & Frith, 2021). Given that the posterior in pre-disambiguation trials depends partially on the distribution of the sensory input, we therefore predicted that guesses made in a low precision environment, i.e., images that induce high levels of ambiguity, would on average decrease the effect of confidence on semantic distance. Visual ambiguity was operationalised as Shannon’s entropy calculated over all participant-provided labels for a specific image at pre-disambiguation (Kwisthout et al., 2017; Shannon, 1948; see Methods). Low entropy therefore reflects that participants responded to a given image using a small and consistent set of labels, likely because the image induced a shared (low ambiguous) interpretation (Van de Cruys et al., 2021).

We first median-split the dataframe from the original study based on the images’ entropy, resulting in two types of “environments”, one with high entropy (i.e., less precise) and one with low entropy images (i.e., more precise). In the low entropy environment (Fig. 4a), the interaction between semantic distance and confidence remained significant (χ^2^_(1)_ = 6.01, p = 0.014, B = 0.04, 95% CI [0.01, 0.07]). Considering the full range of both regressors, confidence had a significant negative slope for low semantic distance guesses, although this effect did not completely reverse within the observed values of semantic distance (see dashed vertical lines, Johnson-Neyman interval [0.04, 1.62]; FDR-corrected interval [0.03, 2.41]; Fig. 4a). The interaction was specifically driven by significant positive slopes of semantic distance for high (b = 0.74, t_(1772)_ = 4.43, p < 0.001, 95% CI [0.41, 1.07]) and mean levels of confidence (b = 0.50, t_(1772)_ = 3.02, p < 0.001, 95% CI [0.18, 0.83]), while semantic distance was not significantly related to Aha! scores for low confident guesses (b = 0.27, t_(1772)_ = 1.24, p = 0.21, 95% CI [−0.15, 0.69]). However, as predicted, in the high entropy environment (Fig. 4b), the interaction model was no longer significant (χ^2^_(1)_ = 2.56, p = 0.108, B = 0.02, 95% CI [−0.01, 0.05]), revealing that the precision of the context impacts insight experiences in our task. We additionally explored the effect of the temporal distance between the Mooney and its corresponding clear version (as an alternative metric of sensory uncertainty), and found that higher temporal distance (i.e., less precise predictions at the timepoint of disambiguation) attenuates the interaction effect as well (see Supplementary Fig. 1). Altogether, these results suggest that uncertain sensory environments weaken the effect of predictions on subsequent experiences of insight.

**Figure 4.**
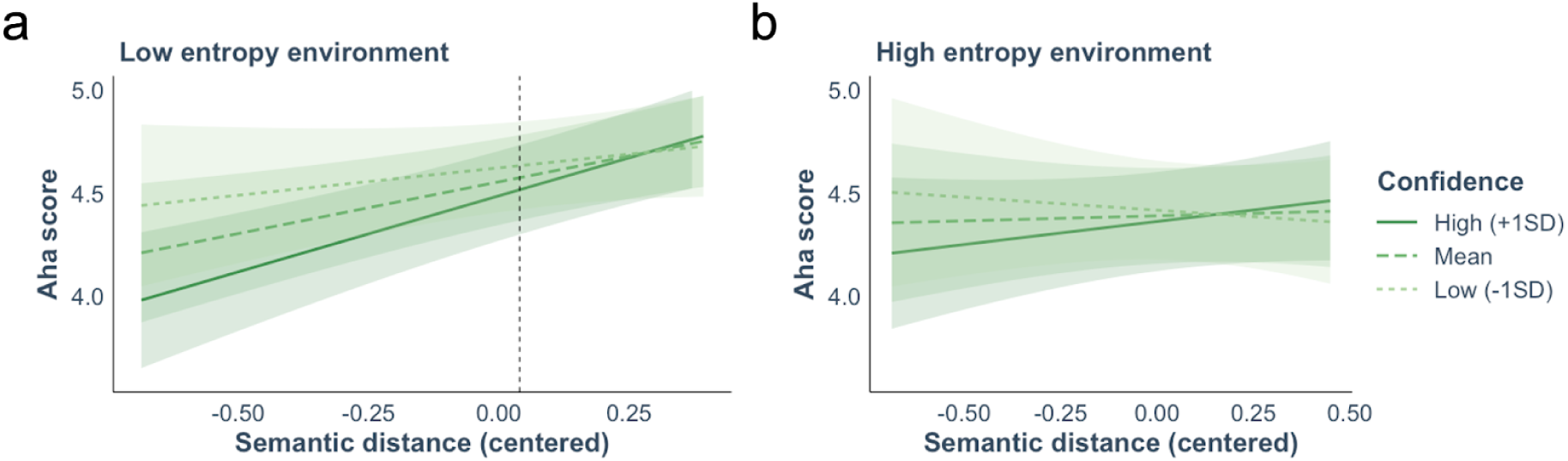
The initial guess has a weaker influence on the insight experience in low precision environments. **a**. In the ‘low entropy environment’, higher confidence reduced insight intensities for low semantic distance guesses. Shaded areas represent 95% confidence intervals, and the dashed vertical line denotes the Johnson-Neyman boundary (uncorrected) below which confidence has a significant slope. **b**. In contrast, there was no significant interaction between semantic distance and confidence in ‘high entropy environments’. Shaded areas represent 95% confidence intervals.

## Discussion

Insight arises when a prediction error is suddenly reduced after a stage of impasse and is often marked by a subjective experience of “Aha!”. Recent proposals have related the intensity of such experience to the magnitude of this prediction error (Becker & Cabeza, 2025; Laukkonen et al., 2023), mostly considering the accuracy of the prediction in explaining Aha! (Becker et al., 2024; Dubey et al., 2023). Here, we formalize this relationship by revealing uncertainty as one of its key components. Specifically, we show that perceptual insight is influenced not only by the accuracy but also the precision of predictions about the content of ambiguous images. In three independent datasets, we find evidence that guesses made with low confidence (low precision) result in higher Aha! ratings if the guess was fairly correct (low semantic distance). The effect reverses for incorrect guesses (high semantic distance): high confidence predictions lead to stronger Aha! experiences than low confidence ones. This pattern was recovered when estimating trial-by-trial prediction errors from participant-provided confidence ratings and the semantic distance between their guess and the target label.

As reported in previous studies (Becker et al., 2024; Dubey et al., 2023; Savinova & Korovkin, 2022; Van de Cruys et al., 2021), we found a positive relationship between the accuracy of predictions, operationalised as semantic distance, and the subjective intensity of insight experiences. Thus, the further away the initial guess from the image’s actual label, the stronger was the reported Aha! experience. However, and importantly so, additionally considering uncertainty of the prediction, here estimated through participants’ confidence ratings, revealed a significant interaction effect. Bayesian inference accounts (and their mathematical implementations) predict a different effect of predictions dependent on the precision awarded to them. For instance, the violation of a weak prediction is less surprising than that of a strong one (Press et al., 2020). The present study corroborated this idea, showing that guesses that were made with higher confidence resulted in less intense insight experiences when the guess was close to the correct label, but in stronger insights when the guess was far from the correct label. This finding highlights the importance of considering both the accuracy and precision of predictions to fully characterise perceptual insight. Moreover, we find that other sources of uncertainty, e.g., the ambiguity (entropy) the Mooney image induces at pre-disambiguation, influenced this relationship. Specifically, the interaction between semantic distance and confidence on the Aha! experience was attenuated in low precision environments, i.e., when the Mooney image induced a high entropy, again highlighting the important role of precision in insight experiences.

We replicated this finding in two additional independent datasets, both showing stronger interaction effects than the original study. For instance, in the “Catch-Online” dataset, the interaction was fully crossed and outweighed the main effect of semantic distance. One likely contributor is the proportion of images that could not be easily disambiguated (i.e., catch images). We speculate that this proportion is itself another determinant of context uncertainty, impacting prediction error signals and, as a consequence, the intensities of insight experiences in the different experiments. Whereas the original study included no catch trials and thus produced uniformly high Aha! scores, the “Catch-Lab” (25% catch) and “Catch-Online” (50% catch) datasets introduced trials that typically do not elicit strong insight experiences, thereby increasing overall variance in Aha! ratings. This broader baseline may have enabled participants to differentiate their insight experiences more precisely. Variance was likely further enhanced by the use of visual analogue scales rather than a 6-point Likert scale, potentially allowing the interaction effect to be captured more sensitively. From a mechanistic perspective, reactivation of the pre-disambiguation posterior may be facilitated when fewer posteriors must be maintained (in studies with higher catch trial proportion), as posteriors for irrelevant catch clear images can be discarded. Under these conditions, even low-confidence posteriors may be sufficiently reactivated to generate a strong prediction error. These successful replications indicate that the effect generalizes robustly across different setups and conditions and is not purely dependent on a mere repetition effect, as no significant interaction was observed in catch trials.

As prediction errors are influenced by the prediction accuracy and precision, the interaction between these two components in explaining the Aha! provides preliminary evidence that insight experiences are associated with prediction errors. Using Kullback-Leibler Divergence, a known formula to quantify the divergence between the prediction and the posterior distribution after observing the data (Itti & Baldi, 2009), we therefore estimated the prediction errors that enable insight experiences, incorporating subjects’ guesses and their reported confidence. Mooney images present an optimal case for studying the content of internal predictions and the influence of the precision awarded to it as the prediction does not immediately reduce all prediction errors at pre-disambiguation. Instead, participants form an idea of what the content of the image reflects based on the available imprecise evidence (the posterior). We exploited the inferential nature of Bayesian inference (Lee & Mumford, 2003; Ma et al., 2023; Wacongne et al., 2011) to propose that this posterior is reactivated during clear image viewing and results in a strong prediction error when convolved with the precise evidence. Prediction errors derived in that way positively predicted Aha! ratings on a trial-by-trial level and largely paralleled empirical results, i.e., guesses made with lower confidence resulted in weaker prediction errors (and less intense insight). The results suggest that experiences of insight seem to read out the magnitude of the prediction error and further strengthen proposals of their relationship (Becker & Cabeza, 2025; Laukkonen et al., 2023). Importantly, this readout likely resembles a second order inference about the sudden reduction of the prediction error, that is awarded a lot of precision, and not the mere content of the prediction error (Laukkonen et al., 2023).

Finally, we confirmed that Aha! ratings measured in our task reproduced the previously reported positive link with subsequent (recognition and resolution) memory. While other studies reported this link for Aha! scores during post-disambiguation (Becker et al., 2025; Van de Cruys et al., 2021), we showed this effect for Aha! recorded during clear image viewing. Together with the on average higher insight intensities during this stage, we propose that Mooney image disambiguation in our task likely arose while being presented with the clear image. These insight-related mnemonic effects are in line with a body of evidence showing that prediction errors are linearly related to better subsequent memory (e.g., Greve et al., 2017), mediated by novelty and surprise as important factors of learning, consistent with predictive-coding models of memory (Friston, 2010; Henson & Gagnepain, 2010). Thus, prediction errors and Aha! experiences produce similar consequences on behaviour, further strengthening their relationship.

The present study investigates the effect of semantic prediction (errors) on the Aha! experience, by considering the semantic dissimilarity between participants’ guesses and the target label of that image. Intuitively, the perceptual dissimilarity should play a key role as well. For instance, if the image showed a banana, guessing it is an apple would be closer semantically, but guessing it is a curved stick could be closer perceptually. These two predictions can in principle decouple and their (interactive) effects on the Aha! experience should be explored in the future. Furthermore, the current study employed a single scale to measure the subjective experience of insight instead of several finer-grained subscales for different facets (e.g., happiness, surprise, suddenness; Danek et al., 2014a). Including finer-grained subscales in future research may help to dissociate subsequent memory effects of more surprise- and positive affect-related components of insight.

Finally, future research should further try to elucidate the underlying mechanisms of the two potentially dissociable axes that lead to subjective insight experiences and whether they have different effects on learning and memory. Specifically, our study indicated that stronger feelings of insight arise when people either make a highly confident but incorrect guess, or a fairly correct but low confident guess. While the first scenario includes a complete update of the previous generative model, the second scenario merely affords an increase in the model’s confidence. In principle, one could predict that the first scenario is more surprising on average and therefore facilitates learning more strongly. Paralleling dual-mechanism ideas on how prediction errors influence subsequent memory (Van Kesteren et al., 2012), one could also predict different mechanisms to underlie the two sides of the Aha! experiences. We propose that computational models could help elucidate this potential distinction. While we have shown here that the Kullback-Leibler Divergence can be used to compute prediction errors in insight tasks, and that the derived errors predict Aha! experiences, we find this opens exciting possibilities to build a full computational model of perceptual insight using Kullback-Leibler Divergence. Specifically, the model should incorporate an inferential process of prediction error minimization at every viewing stage, and a reactivation of the former posterior as the new prior in the subsequent stage. Lastly, we want to highlight that further research is needed to better understand how exactly the greyscale and the Mooney image are associated with each other and how the prediction errors are resolved at this point.

In sum, our results contribute to a recent line of research considering subjective experiences of insight as a readout of prediction errors, but critically highlight the importance to consider the uncertainty about predictions to fully characterize this relationship. This characterization might further facilitate understanding the effect of insight on subsequent memory, which has been sketched to be beneficial, but the mechanisms are less understood.

## Methods

### Participants

A total of 150 native English speakers aged 18 to 40, with approval rates above 95% and at least 10 previous submissions, were recruited online via Prolific. Participants were excluded if they did not complete the study successfully (e.g., due to technical issues; 6 participants) or exhibited no variation across trials in their confidence ratings (pre-disambiguation) or Aha! ratings (greyscale and post-disambiguation). Subsequently, the 1.5 interquartile range (IQR) rule was applied to exclude participants that reported mean identification of (clear) greyscale images or mean recognition memory below the lower bound (< Q1 - 1.5 IQR). Sample size and exclusion criteria were preregistered (https://aspredicted.org/jvf2-r328.pdf, 21.03.2024). After these exclusions, the final sample consisted of 122 participants (50 female, 72 male; M_age_ = 31.36, SD = 5.54; self-reported). Participants were compensated at a rate of £6.36 per hour for approximately 60 minutes of participation. Participation was voluntary, and all participants provided informed consent prior to the study. Ethical approval was obtained from the University of Granada Ethics Committee.

### Stimuli

Stimuli were taken from the THINGS-Mooney database (Linde-Domingo et al., 2024) which is built on the object database THINGSplus (Stoinski et al., 2023). THINGS-Mooney contains 1,854 greyscale transformed object images of concrete picturable and nameable nouns embedded in scenes and their thresholded and Gaussian filtered two-tone versions (‘Mooney images’; for more information, see Linde-Domingo et al., 2025). The database additionally provides metadata for each image per viewing stage, including mean identification scores and semantic distances between participant provided labels in a naming task and the correct object label, based on more than 100,000 ratings.

For the present study, we selected 48 Mooney images that exhibited the greatest change in semantic distance from pre- to post-disambiguation in the database. This was done to ensure a low spontaneous disambiguation rate, i.e., disambiguation before seeing the clear version, and to robustly induce a strong Aha! experience (which we expected to be driven by a large reduction in semantic distance). Additionally, this approach allowed us to infer subject-level effects of resolving visual ambiguity, even though the images were similar in terms of semantic distance. We selected 24 images depicting natural objects and 24 artificial objects. There was no significant difference in pre-disambiguation semantic distance between the categories in the stimulus set (t_(46)_ = 1.09, p = 0.28, d = 0.32, 95% CI [−0.01, 0.04]), suggesting similar baseline difficulty across images. Lastly, all image filenames were renamed into random 3-digit numbers.

For the memory task, we selected 48 lure (*‘new*’) Mooney images (50% natural objects) that matched the to-be-encoded (‘*old*’) Mooney images in pre-disambiguation semantic distance (p = 1) and showed no significant category difference (t_(46)_ = 0.64, p = 0.527, d = 0.18, 95% CI [−0.02, 0.03]). To reduce task duration, participants were tested on half of the encoded images, randomly selected from one out of four previously created subsets. Each subset contained 24 old and 24 new images, balanced so that 50% depicted artificial objects and the other half natural objects. The old and new images in each set were matched in pre-disambiguation semantic distance to ensure consistent identification difficulty.

### Experimental Procedure

The study consisted of two phases: an incidental encoding phase during which participants learned to identify the central object in a Mooney image, and a surprise memory test shortly after the encoding phase. The task was implemented in JavaScript using jsPsych 7.3.4 and hosted on the JATOS server of the Mind, Brain, and Behavior Research Center in Granada, Spain.

In the first phase, participants saw 48 different Mooney images twice, interleaved with the corresponding greyscale version of the image to induce disambiguation (144 trials per subject). Trials were grouped into 12 blocks, with a maximum of 30 seconds break between them if participants did not press a button to continue earlier. In each block, four greyscale images were presented followed by eight Mooney images of which half corresponded to the presented greyscale image (post-disambiguation) and the other half were new images (pre-disambiguation) whose greyscale image would be presented in the next block. Trial order was shuffled randomly for each participant, while keeping the block structure. The maximum distance between each Mooney image and its greyscale version was 10 trials (M_distance pre-grey_ = 4.83 trials, SD = 2.54; M_distance grey-post_ = 4.83 trials, SD = 2.56). Stimuli were presented at a size of 500 x 500 pixels against a white background. The jsPsych *virtual-chinrest* plugin was used to rescale image size to participants’ individual screen setups, i.e. 50 pixels equaled 1 cm.

Each trial started with a fixation cross (500 ms) in the middle of the screen, followed by a Mooney or a greyscale image (4 s) during which participants were instructed to press the spacebar as soon as they identified the presented object, in which case they subsequently rated the intensity of their Aha! experience on a 6-point Likert scale (no Aha - very weak Aha - weak Aha - some Aha - strong Aha - very strong Aha). Note that this restricts Aha! ratings to trials in which participants subjectively identify the image, resulting in relatively more Aha! ratings during grey and post-than pre-disambiguation. Independent of subjective identification and Aha! ratings, all participants were then asked to type the name of the object using the keyboard and to rate their confidence in the provided name on a 6-point Likert scale (guess - unsure - somewhat unsure - somewhat sure - sure - completely sure) before proceeding to the next trial. Participants were instructed to take a guess if they could not identify the image. As participants typed, a drop-down menu matched their response with a predefined list of 3,434 terms, which included the THINGS image names and their WordNet synonyms. This allowed us to use the available semantic embeddings from the THINGS database (Hebart et al., 2019) to calculate the semantic distance between each guess and the correct image label (see Methods “Semantic distance”). All tasks, except for the subjective identification report, were self-paced.

In the second phase, participants’ incidental memory for the Mooney images was assessed using two tasks: a direct recognition memory task and an indirect solution memory task (e.g., Kizilirmak et al., 2016). Participants evaluated 24 old Mooney images from the encoding phase and 24 new images, rating their recognition memory (sure new - probably new - guess new - guess old - probably old - sure old) and whether they identify the Mooney image (sure no - probably no - I don’t think so - I think so - probably yes - sure yes) on a 6-point Likert scale. To reduce order effects between the memory tasks (cf. Kizilirmak et al., 2016), task order was randomized for each trial. Each Mooney image was displayed for 1 second, preceded by a 250-ms fixation cross in the center of the screen. One of the four stimuli subsets was randomly selected for each participant at the beginning of the experiment and the trial order was randomly shuffled to minimize temporal dependence between the encoding and retrieval sequence. Rank correlation between image order in the encoding and the memory phase was tested prior to launching the experiment with 150 simulated datasets (mean p-value = 0.49, 7 significant correlations). Furthermore, the direction of the memory Likert scales was randomly determined for each participant at the start of the experiment and remained constant throughout.

A literature-based definition of the Aha! was provided in the instructions to ensure a common understanding (Becker et al., 2021; Danek et al., 2014a). This definition stated that “The Aha! experience (Eureka effect, insight) often accompanies the sudden switch from not understanding to understanding […] (akin to a light bulb turning on). It is characterized by feelings of happiness, relief, and surprise”. We purposefully did not include confidence as a criterion of the Aha! moment to not confound our confidence and Aha! measure, as mentioned in previous studies (Danek & Salvi, 2020). The 6-point Likert scale was used as recent studies provided evidence that the Aha! is a continuous phenomenon showing intermediate levels of Aha! (Danek & Wiley, 2017, 2020; Webb et al., 2016). We acknowledge that our global Aha! measure cannot separately capture the different facets of the Aha! but due to time constraints in our online experiment we followed the recommendations for likert scales in this type of tasks (Ishikawa et al., 2019a, 2019b; Simms et al., 2019).

### Semantic distance and semantic entropy

To calculate the semantic distance between each participant provided label and the correct label of that image, we used the THINGS database semantic embeddings (Hebart et al., 2019) which provide an estimate of the semantic proximity (or distance) between word meanings based on their co-occurrence statistics in large text corpora. For each concept, a word vector was extracted from word2vec and augmented with WordNet synsets. Semantic distance was then computed as the cosine dissimilarity between the provided and the true embedding ([0, 1]). Note that WordNet synonyms were treated as correct guesses, i.e., cosine dissimilarity was zero.

Semantic entropy over participants’ guesses was computed using Shannon’s formula as in previous work, albeit with more conservative thresholds (Van de Cruys et al., 2021). First, a list of guesses for each image in each viewing stage (e.g., image A at pre-disambiguation) was created. Using fuzzy matching, each guess was compared to the other guesses in the list. If the guess was not yet in the list, a new list entry was initialized - otherwise, the frequency of that guess was increased by one. From these frequencies, we computed the probability of a specific guess made per image per condition, which was then fed into Shannon’s entropy formula (Shannon, 1948). This gave us an information theory estimation of an image’s crowdsourced ambiguity per condition which serves as a proxy of the uncertainty in the visual input. Please note that, unlike previous studies (Van de Cruys et al., 2021), we did not assume post-disambiguation entropy to be necessarily zero. After cleaning the data, we therefore computed the entropy over the guesses at each exposure stage and created two separate dataframes with all guesses, disregarding whether participants identified the Mooney image correctly at post-disambiguation, and another dataframe that included only correct post-disambiguation trials (semantic distance post-disambiguation was 0). We primarily used the latter dataframe for analysis, as the Aha! experience is defined as the experience following a correct solution (Kizilirmak & Becker, 2022).

### Statistical analysis

Confirmatory analyses were pre-registered. Data preprocessing and descriptive analyses were implemented in Python (v 3.11.6). To test the effect of viewing stage, we computed rmANOVAs with Greenhouse-Geisser corrected p-values where applicable and partial eta squared as effect size using Pingouin (v 0.5.4). One-tailed pairwise post-hoc tests were performed using paired t-tests with FDR correction.

Next, we used (linear) mixed models to (i) test the effect of semantic distance and its interaction with confidence on the Aha! intensity including correctly disambiguated trials only (post-disambiguation semantic distance = 0) and (ii) to explore the effect of Aha! and semantic distance on subsequent memory (additionally excluding spontaneous disambiguation trials). Note that the Aha! experience could theoretically arise during clear or post-disambiguation image viewing, depending on when in time the Mooney image can be associated with its corresponding clear image. A significant difference in the distributions of Aha! ratings (χ^2^ = 1997.27, p < 0.001) showed higher Aha! ratings during clear (M = 4.37, SD = 1.62) compared to post-disambiguation viewing (M = 4.07, SD = 1.75) and indicated that a substantial amount of perceptual insights already occurred in clear, greyscale trials. In all analyses, we thus used the Aha! rating in clear trials as dependent variable. Subjective identification pre-disambiguation varied systematically with semantic distance and confidence and was thus entered as a covariate where appropriate (see Supplementary Material).

Mixed model analyses were conducted in R (v 2025.09.2) with the lmer function (v 1.1.35; Bates et al., 2015). Level 1 continuous predictors were mean-centered to the participant before being entered into the model. Level 2 predictors (image-level entropy) were grand-mean centered. Models were built in a top-down fashion, starting with the most complex random effects structure including a random intercept for every participant and random slopes for the variables of interest as well as a random image intercept where applicable (Barr et al., 2013). Random effects structure was simplified until the model reached convergence. Inference statistics (χ^2^) were implemented by comparing each model containing the variables of interest with a corresponding baseline model using likelihood ratio tests. If the full model included an interaction, the baseline model included both independent variables. We report standardized beta (B) estimates and 95% confidence intervals of the independent variable’s slope from the full model. Interaction effects were probed and plotted (model-predicted values) with the default interact plot, simple slope (unstandardized beta b) and Johnson-Neyman functions of the interactions package (v 1.1.5; Long, 2019) that allow for a more continuous characterization of the interaction between two continuous predictors (Bauer & Curran, 2005; Ji, 2016; Lin, 2020). To account for multiple testing, FDR-correction was applied to the Jonson-Neyman interval. All other p-values (one-sided) are uncorrected unless otherwise noted. Low levels of collinearity were confirmed for all models, and normality of model residuals was checked visually.

For simplicity and increased interpretability, we report linear mixed models in the main manuscript, following recent suggestions that parametric models are robust enough to approximate ordinal data if sample size is large enough and the ordinal variable has sufficient categories (Kromrey & Rendina-Gobioff, 2002; Robitzsch, 2020; Sullivan & Artino, 2013; Taylor et al., 2006). However, we additionally confirmed all models with cumulative link mixed models for ordinal data with the clmm function from the ordinal package (v 2023.12.4.1; Christensen, 2010; see Supplementary Material “Cumulative Link Mixed Models”). Note that in the “Catch-Lab” and “Catch-Online” datasets, Aha! was rated on a visual analogue scale, suitable for analysis with linear mixed models.

Please note that we had pre-registered to combine semantic distance and confidence into one rating to test their interactive effect on Aha!, but we reasoned the current approach, i.e., including both variables and their interaction term in the same model, was more straightforward and did not involve artificially defining at which value the sign of the confidence effect switched. Additionally, subsequent memory analyses were planned to be run on the binary outcome, but as memory performance for old items was at ceiling (M_recognition_ = 0.93, SD = 0.25; M_resolution_ = 0.90, SD = 0.30), we opted to run linear mixed models to capture small differences in memory performance. For that same reason, we did not run comparative ROC analyses to disentangle the contribution of familiarity and recollection to retrieval.

### Kullback-Leibler Divergence

To mathematically derive prediction errors from participant’s semantic distance to the correct label and their confidence in that label, we first scaled all variables between 0 and 1. As before, only correctly disambiguated trials were analysed. At pre-disambiguation, participants encounter an ambiguous input (imprecise evidence) without a concrete prior (flat prior). Thus, their posterior (prior * evidence) closely resembles the ambiguous input. We assume that this posterior will largely reflect participants’ response during pre-disambiguation trials, thus the posterior’s mean tendency and the variance can be described by the semantic distance of participant’s verbal label and participant’s confidence rating, respectively. Subsequently, this posterior is reactivated during greyscale image presentation, acting as a new prior that is convolved with the highly precise sensory evidence of the clear image, leading to a large model update (the greyscale posterior). For simplicity, we assume that the mean tendency of the likelihood function at disambiguation (greyscale viewing stage) is 0 with a variance of 0.1, such that the distance in mean tendency between the prior and likelihood function reflects the semantic distance from a correct label. The prediction error during greyscale viewing is calculated using Kullback-Leibler Divergence between the prior and the posterior, capturing the non-overlap between the distributions, i.e. the amount of surprise due to a change in predictions before (prior) and after the sensory evidence has been processed (posterior; Press et al., 2020). In a final step, the prediction error for every trial is convolved with an error term sampled from a Gaussian distribution to imitate distortions of participant’s readout and converted into a discrete scale that mimics the range of the Aha!.

### “Catch-Lab” Dataset

A total of 32 Spanish Natives were recruited to take part in an in-person study at the University of Granada. Participants were compensated with course credits or the corresponding monetary compensation. Participation was entirely voluntary and all participants provided informed consent. Ethical approval was obtained from the University of Granada Ethics Committee. After exclusion using the same criteria as explained above (with exception of the mean recognition scores, as participants did not have a retrieval phase), a final sample of 30 participants entered the analyses (20 females, 10 male; M_age_ = 21.23, SD = 3.10).

For this dataset, we selected 64 Mooney images based on the same criteria as before (highest change in semantic distance from pre- to post-disambiguation, 50% artificial images, no category difference in pre-disambiguation semantic distance). Importantly, this study included 25% catch trials, i.e., for 16 of the 64 Mooney images we did not present the correct greyscale image. Instead, we showed a different greyscale image from the opposite category with a minimal difference in semantic distance to the correct greyscale image. Catch images were counterbalanced across participants such as that each image was selected the same amount of times as a catch image.

First, participants viewed all 64 Mooney images in a random order for 1500 ms preceded by a fixation period of 1500 ms. Participants’ task was to name the images and to provide a confidence rating on a visual analogue scale (not sure at all - very sure). Then, all 64 Mooney images were presented twice, interleaved with either the corresponding (or the catch) greyscale image to induce (or hinder) disambiguation (192 trials per subject). As before, trials were grouped into blocks of four greyscale images. Trial order was shuffled randomly for each participant, while keeping the block structure and under the constraint of maximum 2 catch images per block. Stimuli were presented at 6 visual degrees and against a grey background. Each trial started with a red fixation dot (jitter 1300 - 1700 ms) in the middle of the screen, followed by a Mooney or greyscale image (1500 ms) together with the fixation dot. Afterwards, participants rated their subjective identification on a visual analogue scale (sure no - sure yes) and, if subjective identification was above 50, the intensity of their Aha! on the same visual analogue scale (no Aha - very strong Aha). Again, a literature-based definition of the Aha! was provided in the instructions. Lastly, participants completed the naming task for all 64 Mooney images. In contrast to the original study, we want to highlight the different setup (in-lab, Spanish sample), the different procedure (naming - disambiguation task - naming), and the added catch trials (25%).

Because this study was in Spanish, we first translated all of the image labels and their synonyms with the help of a LLM (GPT-3.5). Specifically, we prompted the LLM to translate the word to Spanish from Spain (not from other Spanish-speaking regions). Synonyms were added if there were clear synonyms in Spanish, or if the synonym from the English list had a clear Spanish translation. In the case of homonyms (the same word referring to a different meaning), the translation was compared to the image from the database (Stoinski et al., 2023). Lastly, two trained native Spanish speakers manually confirmed the translations. To calculate semantic distance, we back-translated the selected word to English and computed the cosine dissimilarity between the vector embedding for the participant provided label and the correct label as described above. Semantic distance and confidence were mean-centered to participants. Mixed models were performed as described above, once for correctly disambiguated regular trials and once for catch trials.

### “Catch-Online” Dataset

For this dataset, a total of 120 native English speakers were recruited from Prolific. After exclusion using the criteria described above, a final sample of 107 participants (64 female, 52 male; M_age_ = 28.55, SD = 4.51) entered the analyses. Participants were compensated at a rate of £6.00 per hour for approximately 60 minutes of participation. Participation was entirely voluntary and all participants provided informed consent. Ethical approval was obtained from the University of Granada Ethics Committee. This time, 32 Mooney images were selected with the same criteria, as well as 32 greyscale catch images from the opposite category. Every participant saw the correct greyscale image for half of the Mooney images (50% catch trials). The assignment of images to the conditions was again counterbalanced across participants.

During encoding, participants saw all 32 Mooney images twice, interleaved with the (regular or catch) greyscale images (96 trials per subject). As before, trials were grouped into blocks. Each trial started with a fixation cross (1000 ms) in the middle of the screen, followed by the Mooney or greyscale image (2000 ms) at a size of 303 pixels. Afterwards, participants rated their subjective identification on a visual analogue scale (sure no - sure yes), their Aha! (if subjectively identified), and provided a label as well as their confidence (visual analogue scale) in that label. Participants then completed a retrieval phase. Semantic distance was calculated in the same way as described before, using the available embeddings. Semantic distance and confidence were again mean-centered to participants and regular, correctly identified trials and catch trials were analysed separately. Subjective identification was included as a variable of no interest. In contrast to the original and the “Catch-Lab” dataset, this dataset contained 50% catch trials.

## Data availability

The data that support the findings of this study are available at https://osf.io/uyst6.

## Code availability

The code to reproduce the results of this study is available at https://osf.io/uyst6.

## Acknowledgments

J.V. was supported by an FPI grant (CEX2021-001161-M-20-5) by the Spanish Ministry of Science and Innovation. J.L.D. was supported by Project PID2023-151104NA-I00 funded by MCIN/AEI/10.13039/501100011033 and by FEDER, EU, and Grant RYC2021-033940-I funded by MCIN/AEI/10.13039/501100011033 and by the European Union NextGeneration EU/PRTR. C.G.G. was supported by Project PID2023-149428NB-I00 funded by MCIN/AEI/10.13039/501100011033 and by FEDER, EU, and Grant RYC2021-033536-I funded by MCIN/AEI/10.13039/501100011033 and by the European Union NextGeneration EU/PRTR. The Mind, Brain and Behavior Research Center receives funding from grants CEX2023-001312-M by MICIU/AEI/10.13039/501100011033 and UCE-PP2023-11 by the University of Granada. The funders had no role in the study design, data collection and analysis, decision to publish or preparation of the manuscript. This manuscript is part of the PhD thesis of J.V. We thank Clare Press for her valuable feedback on previous drafts of the manuscript.

## Author contributions

All authors were involved in conceptualization, investigation, methodology, project administration, and writing of the original draft and further edits. J.V. was involved in data curation, formal analysis and visualization. C.G.G. and J.L.D. were responsible for funding acquisition, resources and supervision.

