## Supplementary Material for "Bayesian surprise tracks the strength of perceptual insight"

Johannah Völler
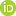
<https://orcid.org/0009-0008-0500-7494>^1,2^✉, Juan Linde-Domingo
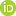
<http://orcid.org/0000-0002-3301-7453>^1,2^*, and Carlos González-García
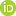
<https://orcid.org/0000-0001-6627-5777>^1,2^* ✉

^1^Mind, Brain and Behavior Research Center, University of Granada, Granada, Spain
^2^Department of Experimental Psychology, University of Granada, Granada, Spain

*Both authors contributed equally

### Supplementary Material

### Linear Mixed Models

#### Subjective identification as covariate

Subjective identification was included as a covariate of no interest in all models where applicable, as it varied systematically with semantic distance and confidence. Specifically, subjective identification negatively predicted semantic distance, such as that semantic distance was lower for higher subjective identification scores (B = -0.17, 95% CI [-0.20, -0.14], t_(3844)_ = -10.85, p < .001). Additionally, subjective identification positively predicted confidence ratings, as higher confidence ratings were observed when participants affirmed subjectively identifying the image (B = 0.41, 95% CI [0.38, 0.44], t_(3859)_ = 28.00, p < .001).

#### Low precision environments attenuate the effect of confidence - Temporal Distance

Previous studies explored the role of the temporal distance between Mooney and their corresponding greyscale images on subjective identification and found no effect, i.e., greyscale images that were presented closer in time to the Mooney images did not result in higher identification rates (Chang et al., 2016). However, no insight ratings were recorded in that study. As the feeling of insight is proposed to arise from a reduction in prediction error, and in order to experience that prediction error one must reactivate one’s belief about the image’s content from pre-disambiguation, we reasoned that the initial prediction should have a stronger effect on the intensity of insight if the guess was temporally closer to the point in time the image can be disambiguated, i.e., when the clear solution image is shown. In other words, initial predictions might be more easily reactivated the closer in time they occurred to the solution image and are therefore more precisely represented, exerting a stronger influence on the insight experience, independent of the ability to disambiguate the image.

To test this, we median-split the dataframe based on this temporal distance and reran the mixed model analysis including the interaction effect of semantic distance and confidence. These led to two groups of trials, one with low and another with high temporal distance. In both groups of trials, the interaction effect remained significant (low temporal distance: χ^2^_(1)_ = 8.68, p = 0.003, B = 0.04, 95% CI [0.01, 0.07]; high temporal distance: χ^2^_(1)_ = 8.21, p = 0.004, B = 0.05, 95% CI [0.02, 0.09]). Crucially, when the temporal distance was small, confidence had a significant negative slope for low semantic distance, and a significant positive slope for high semantic distance guesses (see dashed vertical lines, Johnson-Neyman interval [-0.17, 0.41]; FDR-corrected interval [-0.23, 0.64]; Fig. 1a). Simple slope analyses revealed a significant positive slope of semantic distance for high (b = 0.70, t_(1985)_ = 4.02, p < 0.001, 95% CI [0.36, 1.03]) and mean levels of confidence (b = 0.42, t_(1985)_ = 2.33, p = 0.02, 95% CI [0.07, 0.76]) but no significant slope of low confidence guesses (b = 0.13, t_(1985)_ = 0.59, p = 0.55, 95% CI [-0.31, 0.58]).

In high temporal distance trials, confidence had a significant negative slope for low semantic distance values, but this pattern did not reverse for high semantic distance for the observed values (see dashed vertical lines, Johnson-Neyman interval [0.06, 0.97]; FDR-corrected interval [0.04, 1.18]; Fig. 1b). Additionally, semantic distance had a significant positive slope only for high levels of confidence (b = 0.40, t_(1470)_ = 2.13, p = 0.03, 95% CI [0.03, 0.76]), not for guesses made with mean (b = 0.07, t_(1470)_ = 0.40, p = 0.69, 95% CI [-0.29, 0.44]) or low confidence (b = -0.25, t_(1470)_ = -1.00, p = 0.32, 95% CI [-0.73, 0.23]), suggesting that the sensory uncertainty of the context impacts the prediction error computation in our task.

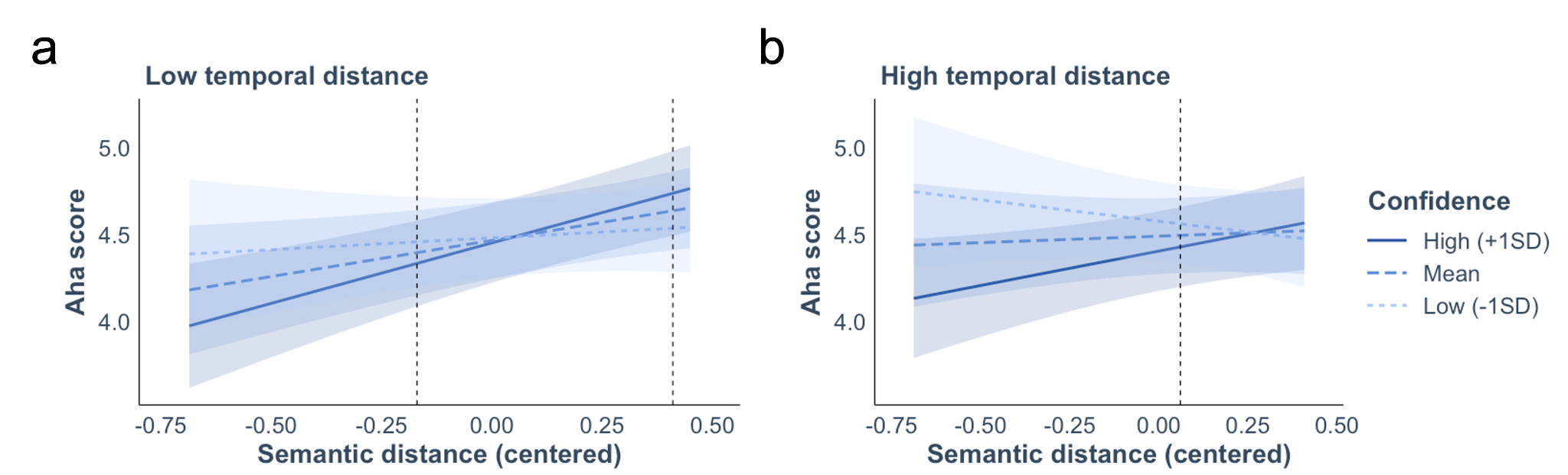

**Figure 1**. Higher temporal distance attenuates the effect of the initial prediction. **a**. In the ‘low temporal distance’ trials, confidence reduced insight intensities for low semantic distance guesses, but increased them for (very) high semantic distance guesses. Shaded areas represent 95% confidence intervals, and dashed vertical lines denote the Johnson-Neyman interval (uncorrected) outside of which confidence has a significant slope. **b**. In contrast, confidence only had a significant negative slope for low semantic distance guesses, while the opposite was not observed for high semantic distance guesses. Shaded areas represent 95% confidence intervals, and dashed vertical lines denote the Johnson-Neyman interval (uncorrected) outside of which confidence has a significant slope.

### Cumulative Link Mixed Models

Cumulative Link Mixed Models, also known as ordered regression models, are suited to analyse ordinal data. Estimation is via maximum likelihood and mixed models are fitted with the Laplace approximation and adaptive Gauss-Hermite quadrature. Models include correct disambiguation trials only and are fitted with a random subject intercept only due to convergence issues for more complex random effects structures. Inferential statistics are implemented with likelihood ratio tests between the model containing the variable of interest and a baseline model. Unstandardized regression estimates are reported for the main effect of interest.

#### Perceptual insight increases memory

The insight models, compared to models without the main effect of insight, predicted ordinal recognition memory strength and subjective identification scores from participant-mean centered Aha! ratings and included a random subject intercept. Paralleling results obtained from linear mixed models, Aha! scores significantly and positively predicted both recognition memory strength (χ^2^_(1)_ = 15.57, p < 0.001, b = 0.29, unstandardized) and subjective resolution (χ^2^_(1)_ = 23.16, p < 0.001, b = 0.31, unstandardized).

#### Semantic distance predicts the intensity of insight experiences

The baseline model predicted ordinal Aha! ratings from binary subjective identification scores and a random subject intercept. The semantic distance mode additionally included semantic distance as the main variable of interest. Model comparison using likelihood ratio tests revealed a main effect of semantic distance (χ^2^_(1)_ = 46.23, p < 0.001, b = 0.94). The effect remained significant when excluding pop-out solution trials < 2 seconds (χ^2^_(1)_ = 15.78, p < 0.001, b = 0.76), paralleling the results obtained with linear mixed models.

#### Confidence moderates the effect of semantic distance on insight

The baseline model predicted ordinal Aha! ratings from binary subjective identification scores, semantic distance, and confidence, and included a random subject intercept. The interaction model additionally included an interaction term between semantic distance and confidence. Model comparison using likelihood ratio tests revealed a significant interaction between semantic distance and confidence on Aha! scores (χ^2^_(1)_ = 18.3, p < 0.001, b = 0.42; Fig. 2). The effect remained significant when excluding pop-out solution trials < 2 seconds (χ^2^_(1)_ = 17.81, p < 0.001, b = 0.53).

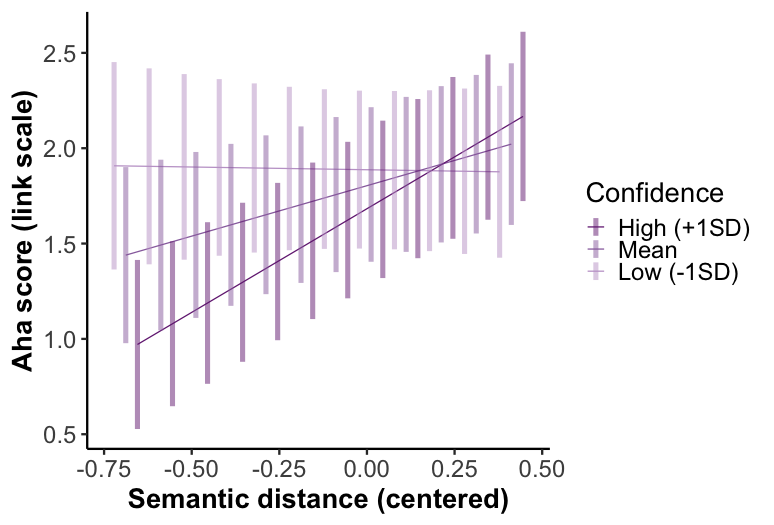

**Figure 2**. Confidence moderates the effect of semantic distance on the intensity of insight (cumulative link mixed models). Model-predicted Aha! values for high (+ 1SD), mean, and low (- 1 SD) levels of confidence indicate that high confidence reduces Aha! ratings for low semantic distance guesses, but increases them for (very) high semantic distance guesses. Shaded areas represent 95% confidence intervals.

#### Low precision environments attenuate the effect of confidence

##### Entropy

The baseline and full model were initialised as explained above after median-splitting the dataframe based on entropy. The interaction effect between semantic distance and confidence was significant in the low entropy environment (χ^2^_(1)_ = 5.22, p = 0.022, b = 0.31; Fig. 3a). In the high entropy environment, the interaction effect was also significant, in contrast to the results obtained from the linear mixed models, but numerically smaller (χ^2^_(1)_ = 4.18, p = 0.041, b = 0.32; Fig. 3b).

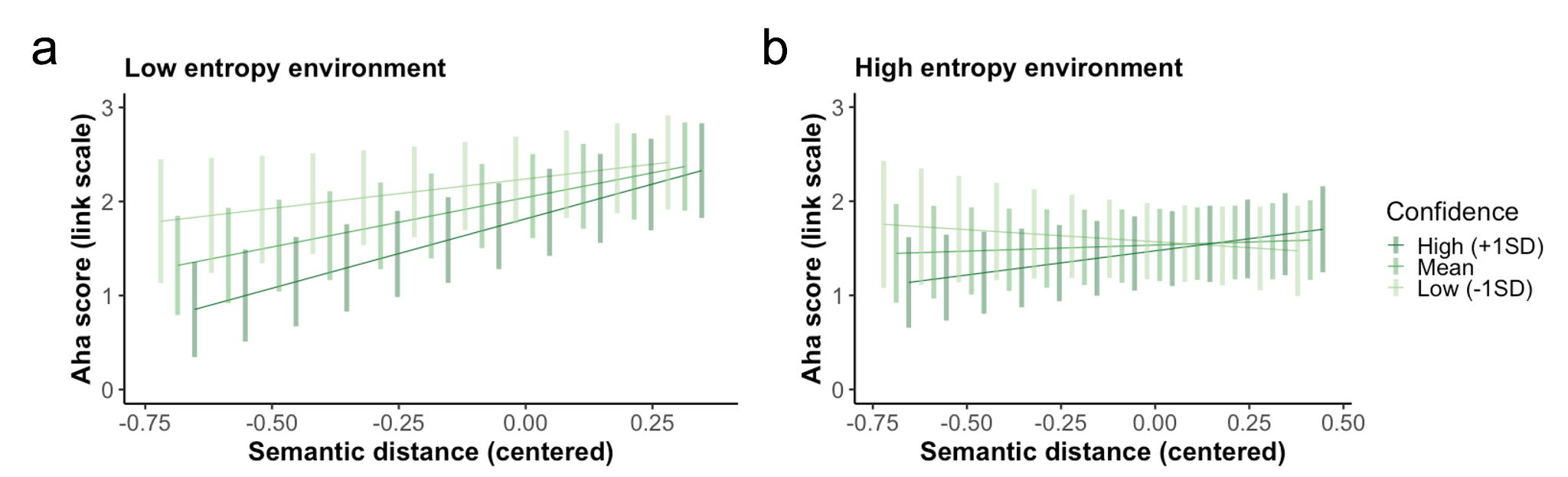

**Figure 3.** The initial guess has a weaker influence on the insight experience in low precision environments (cumulative link mixed models). **a**. In the ‘low entropy environment’, higher confidence reduced insight intensities for low semantic distance guesses. Shaded areas represent 95% confidence intervals. **b**. In contrast, there was no significant interaction between semantic distance and confidence in ‘high entropy environments’. Shaded areas represent 95% confidence intervals.

##### Temporal Distance

The baseline and full model were initialised as explained above. The interaction effect between semantic distance and confidence was significant in the low temporal distance environment (χ^2^_(1)_ = 12.51, p < 0.001, b = 0.47; Fig. 4a) and in the high temporal distance environment (χ^2^_(1)_ = 5.57, p = 0.018, b = 0.35; Fig. 4b), paralleling the results found with linear mixed models.

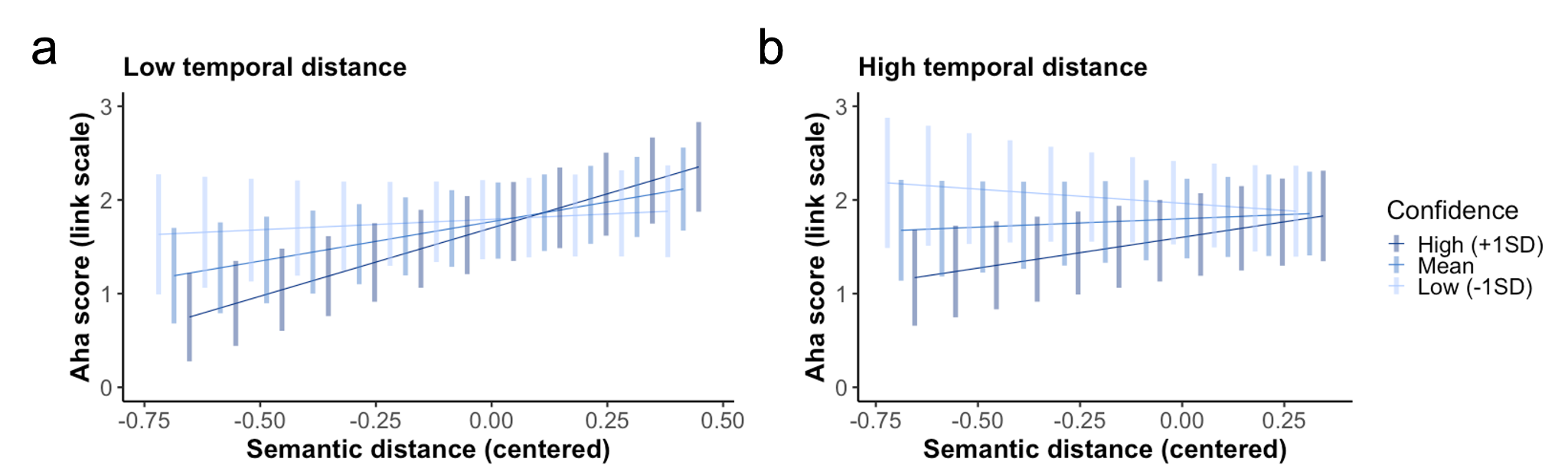

**Figure 4**. Higher temporal distance attenuates the effect of the initial prediction (cumulative link mixed models). **a**. In the ‘low temporal distance’ trials, confidence reduced insight intensities for low semantic distance guesses, but increased them for (very) high semantic distance guesses. Shaded areas represent 95% confidence intervals. **b**. In contrast, confidence only had a significant negative slope for low semantic distance guesses, while the opposite was not observed for high semantic distance guesses. Shaded areas represent 95% confidence intervals.
